# Antimicrobial Resistance and Genomic Characterization of *Salmonella* Dublin Isolates in Cattle from the United States

**DOI:** 10.1101/2021.03.23.436584

**Authors:** Mariela E. Srednik, Kristina Lantz, Jessica A. Hicks, Brenda R. Morningstar-Shaw, Tonya A. Mackie, Linda K. Schlater

## Abstract

*Salmonella enterica* subspecies *enterica* serovar Dublin is a host-adapted serotype in cattle, associated with enteritis and systemic disease. While rare in humans, it can cause severe illness, including bacteremia, with hospitalization and death. In the United States, *S*. Dublin has become one of the most multidrug-resistant serotypes. The objective of this study was to characterize *S*. Dublin isolates from sick cattle by analyzing phenotypic and genotypic antimicrobial resistance (AMR) profiles, the presence of plasmids, and phylogenetic relationships. *S*. Dublin isolates (n=140) were selected from submissions to the NVSL for *Salmonella* serotyping (2014 – 2017) from 21 states. Isolates were tested for susceptibility against 14 class-representative antimicrobial drugs. Resistance profiles were determined using the ABRicate with Resfinder and NCBI databases, AMRFinder and PointFinder. Plasmids were detected using ABRicate with PlasmidFinder. Phylogeny was determined using vSNP. We found 98% of the isolates were resistant to more than 4 antimicrobials. Only 1 isolate was pan-susceptible and had no predicted AMR genes. All *S*. Dublin isolates were susceptible to azithromycin and meropenem. They showed 96% resistance to sulfonamides, 97% to tetracyclines, 95% to aminoglycosides and 85% to betalactams. The most common AMR genes were: sulf2 and tetA (98.6%), aph(3’’)-Ib and aph(6)-Id (96.4%), floR (94.3%), and blaCMY-2 (85.7%). All quinolone resistant isolates presented mutations in *gyr*A. Ten plasmid types were identified among all isolates with IncA/C2, IncX1, and IncFII(S) being the most frequent. The *S*. Dublin isolates show low genomic genetic diversity. This study provided antimicrobial susceptibility and genomic insight into *S*. Dublin clinical isolates from cattle in the U.S. Further sequence analysis integrating food and human origin *S*. Dublin isolates may provide valuable insight on increased virulence observed in humans.

## Introduction

The CDC estimates that each year in the United States, *Salmonella enterica* causes 1.2 million infections, 24,000 hospitalizations, and 450 deaths. According to FoodNet data, *Salmonella* Dublin was more commonly isolated from blood (61%) than were other *Salmonella* (5%) [1]. According to surveillance data from the National Antimicrobial Resistance Monitoring System (NARMS), the proportion of resistant isolates is higher among *S*. Dublin than among other serotypes. A 2019 *S*. Dublin outbreak in the U.S. was linked to ground beef with 13 cases reported with 9 hospitalizations and 1 death in 8 states. Despite the relatively low incidence of human cases, zoonotic or foodborne transmission of *S*. Dublin is of high concern because of the increased antimicrobial resistance and elevated hospitalization and death rate in humans.

Salmonellosis may cause severe disease in cattle and poses a significant zoonotic risk. Farm workers, calf handlers, and their families are clearly at risk of becoming infected by *Salmonella* spp. during outbreaks of clinical illness, but the risk of exposure goes far beyond farm workers or veterinarians with direct animal contact during outbreaks of disease. Asymptomatic shedding of *Salmonella*, a characteristic of *Salmonella* Dublin infection, is also an issue with other common cattle serovars such as Newport and Typhimurium, and creates risk for people in direct contact with the animal, its feces, or milk. There is also risk for foodborne transmission from exposure to contaminated meat from cattle infected with *Salmonella* Dublin, including dairy beef and cull dairy cows, typically via fecal contamination of the carcass at the time of slaughter [2].

In the United States, *Salmonella* Dublin has become one of the most multidrug-resistant (MDR) serotypes. The increasing prevalence of *Salmonella* Dublin infection in the U.*S*. dairy industry and its unique status as host-adapted in cattle merit more specific attention [3]. A National Veterinary Services Laboratories (NVSL) *Salmonella* serotyping report from a 2017 study [4] demonstrated that of the 1,655 *Salmonella* isolates identified at the NVSL from clinical bovine case submissions, the most common serotype was *Salmonella* Dublin (26.4%), followed by *Salmonella* Cerro (17%) and *Salmonella* Montevideo (10.3%).

The antimicrobials of choice for treating bacterial gastroenteritis in humans are generally the fluoroquinolone ciprofloxacin for adults and the cephalosporin ceftriaxone for children.

Antimicrobial drugs are essential to protect animal health in livestock production systems [5]. *Salmonella* Dublin is a host-adapted serovar that can cause significant levels of morbidity and mortality, particularly in dairy calves, potentially necessitating antimicrobial treatment. However, the multidrug resistant nature of *S*. Dublin and limited range of approved antimicrobials often limit treatment options to supportive symptomatic treatment. If antimicrobial treatment is used without testing the susceptibility of the bacteria, treatment may be ineffective and contribute to increasing antimicrobial resistance. Because of the zoonotic implications of this disease, responsible use of antimicrobials in treatment is a critically important aspect of *S*. Dublin management, both for animal and human health.

The mechanism by which *S. enterica* typically develop antimicrobial resistance (AMR) differs according to the drug. Fluoroquinolone resistance typically occurs through clonal dissemination of *Salmonella* isolates with chromosomal mutations conferring resistance, while cephalosporin resistance usually is acquired by acquisition of mobile genetic elements via plasmids and transposons.

The objective of this study was to compare *Salmonella* Dublin isolates from clinical cattle samples throughout the United States during the period 2014-2017 and to analyze antimicrobial resistance profiles, presence of plasmids, and phylogenetic relationships by geographic distribution and period of time.

## Materials and methods

### Bacterial isolates

*Salmonella* Dublin clinical isolates from cattle (n=140) were selected, 110 (78.5%) from infection and 30 (21.4%) unknown, from 2014-2017 submissions for *Salmonella* serotyping archived at the NVSL. Samples came from 21 U.S. States (MN n=37, IA n=21, NY n=17, SD n=10, OH n=7, IL n=6, TX n=6, IN= 5, WA=5, MO n=5, KY=4, ID n=2, WI=2, MD=1, AL=1, NE n=1, KS n=1, UT n=1, MI n=1, FL n=1).

The dataset was initially limited to one sample per year per owner. If more than the targeted number of isolates remained, a randomly selected subset of isolates was chosen. The data was then de-identified to remove information other than the animal species, state of origin, clinical status, and sample type and assigned a unique identifier. Identity was confirmed using Biotyper software with an Autoflex Speed MALDI-TOF instrument (Bruker Daltonics).

### Antimicrobial susceptibility testing

All *Salmonella* isolates were tested for antimicrobial susceptibility against 14 class-representative antimicrobial agents using the Sensititre CMV4AGNF plate (Thermo Scientific), including: gentamicin (GEN), streptomycin (STR), amoxicillin/clavulanic acid (AMC), cefoxitin (FOX), ceftriaxone (CRO), meropenem (MER), sulfisoxazole (FIS), trimethoprim/sulfamethoxazole (SXT), ampicillin (AMP), chloramphenicol (CHL), ciprofloxacin (CIP), nalidixic acid (NAL), azithromycin (AZM), and tetracycline (TET). Interpretation criteria were established by the NARMS.

### Identification of antimicrobial resistant genotype

*Salmonella* Dublin isolates were subjected to whole genome sequencing (WGS) with the Illumina MiSeq platform using 2×250 paired end chemistry and the NexteraXT library preparation kit. AMR gene alleles were detected using AMRFinder [6] and ABRicate [7] with the NCBI and ResFinder databases. Plasmid replicons were identified using the PlasmidFinder [9] database and ABRicate. PointFinder was used for analysis of chromosomal point mutations [10].

### Relationship of antimicrobial susceptibility with antimicrobial genes

Using the phenotypic results as the reference outcome, sensitivity was calculated by dividing the number of isolates that were genotypically resistant by the total number of isolates exhibiting clinical resistance phenotypes. Specificity was calculated by dividing the number of isolates that were genotypically susceptible by the total number of isolates with susceptible phenotypes.

### Phylogenetic analysis

Phylogenetic analysis was performed with vSNP (https://github.com/USDA-VS/vSNP) using the *S*. Dublin accession CP01917.1 as a reference.

## Results

### Antimicrobial susceptibility testing

Of the 140 isolates examined, 138 (98.5%) were resistant (R) to more than 4 antimicrobials. The most common resistance profile was: AMC, AMP, FOX, CRO, CHL, STR, SUL, TET in 109 (77.8%) isolates. Overall, 96% and 97% of the isolates were resistant to FIS and TET, respectively, followed by 95% with aminoglycoside resistance and 85% with beta-lactam resistance (Table 1). All of the *Salmonella* Dublin isolates were susceptible to AZM and MER. One isolate was resistant only to ampicillin presenting only the blaTEM-1 gene. One isolate was pan-susceptible with no predicted AMR genes.

**Table 1.**
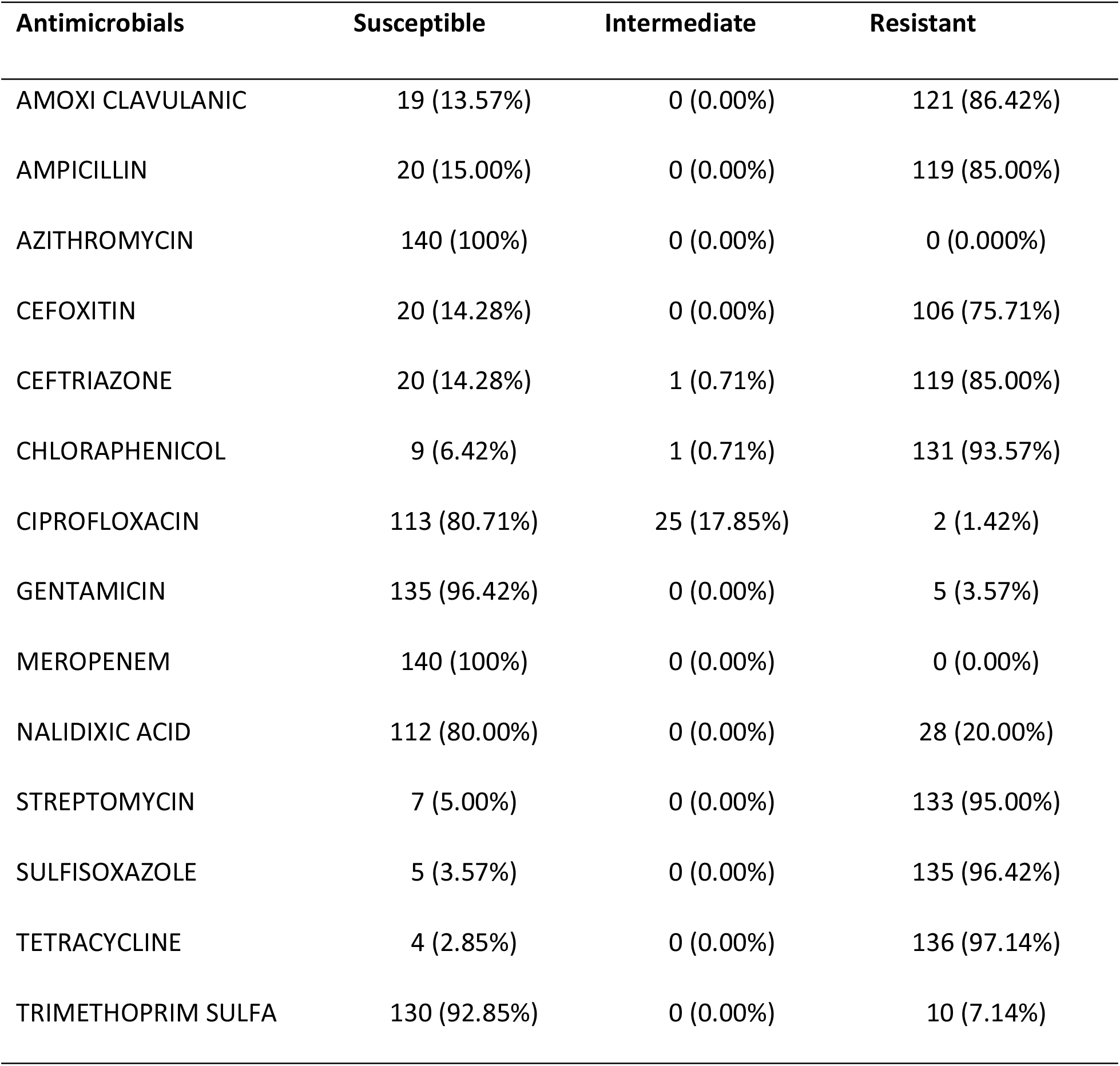
Antimicrobial resistance rates of the 140 *Salmonella* Dublin isolates.

### Antimicrobial resistance genes

The most commonly found antimicrobial resistance genes (ARG) were sulf2 and tetA, conferring resistant to sulfonamides (n=138; 98.57%) and tetracycline (n=138; 98.57%), respectively. Other frequently identified genes were aph(3’’)-Ib and aph(6)-Id genes, both detected in 135 isolates (96.42%). The floR gene was the most commonly identified gene conferring resistant to chloramphenicols (n=132; 94.28%). BlaCMY-2 was also frequently present and confers resistance to all beta-lactams (n=120; 85.71%) (Table 2). Resistance genes for macrolides or meropenem were not found.

**Table 2.** Antimicrobial resistance genes of 140 *Salmonella* Dublin isolated from cattle.

| Drug classes | Resistance genes | Number of isolates | Positive rates (%) |
| --- | --- | --- | --- |
| AMINOGLYCOSIDES | ant(2'')-Ia | 5 | 3.57 |
|  | aph(3'')-Ib | 135 | 96.42 |
|  | aph(6)-Id (strB) | 135 | 96.42 |
|  | aph(3')-Ia | 50 | 35.71 |
| BETA-LACTAMS | blaCMY-2 | 120 | 85.71 |
|  | blaCMY-130 | 1 | 0.71 |
|  | blaCMY-132 | 1 | 0.71 |
|  | blaTEM-1A | 1 | 0.71 |
|  | blaTEM-1B | 67 | 46.85 |
|  | blaTEM-130 | 1 | 0.71 |
| CHLORAMPHENICOLS | floR | 132 | 94.28 |
|  | cmlA5 | 4 | 2.85 |
| QUINOLONES | qnrB19 | 2 | 1.42 |
| SULFONAMIDES | sul1 | 3 | 2.14 |
|  | sul2 | 138 | 98.57 |
| TETRACYCLINES | tetA | 138 | 98.57 |

In addition, two ARGs were found only in the Resfinder database but not in the NCBI/AMRFinder. These genes, mdfA and aac(6’)-Iaa, were detected in all isolates using the Resfinder database (n=140; 100%).

The most frequent AMR gene profile was: blaCMY-2, floR, aph(3’’)-Ib, aph(6)-Id, sul2, tetA in 52 (37.1%) isolates.

### Point mutations

Some isolates (n=25, 17.85%) exhibited weak drug resistance (intermediate) to ciprofloxacin; and they presented chromosomal structural gene mutations in the *gyr*A gene (Fig 1). For nalidixic acid, 28

**Fig 1.**
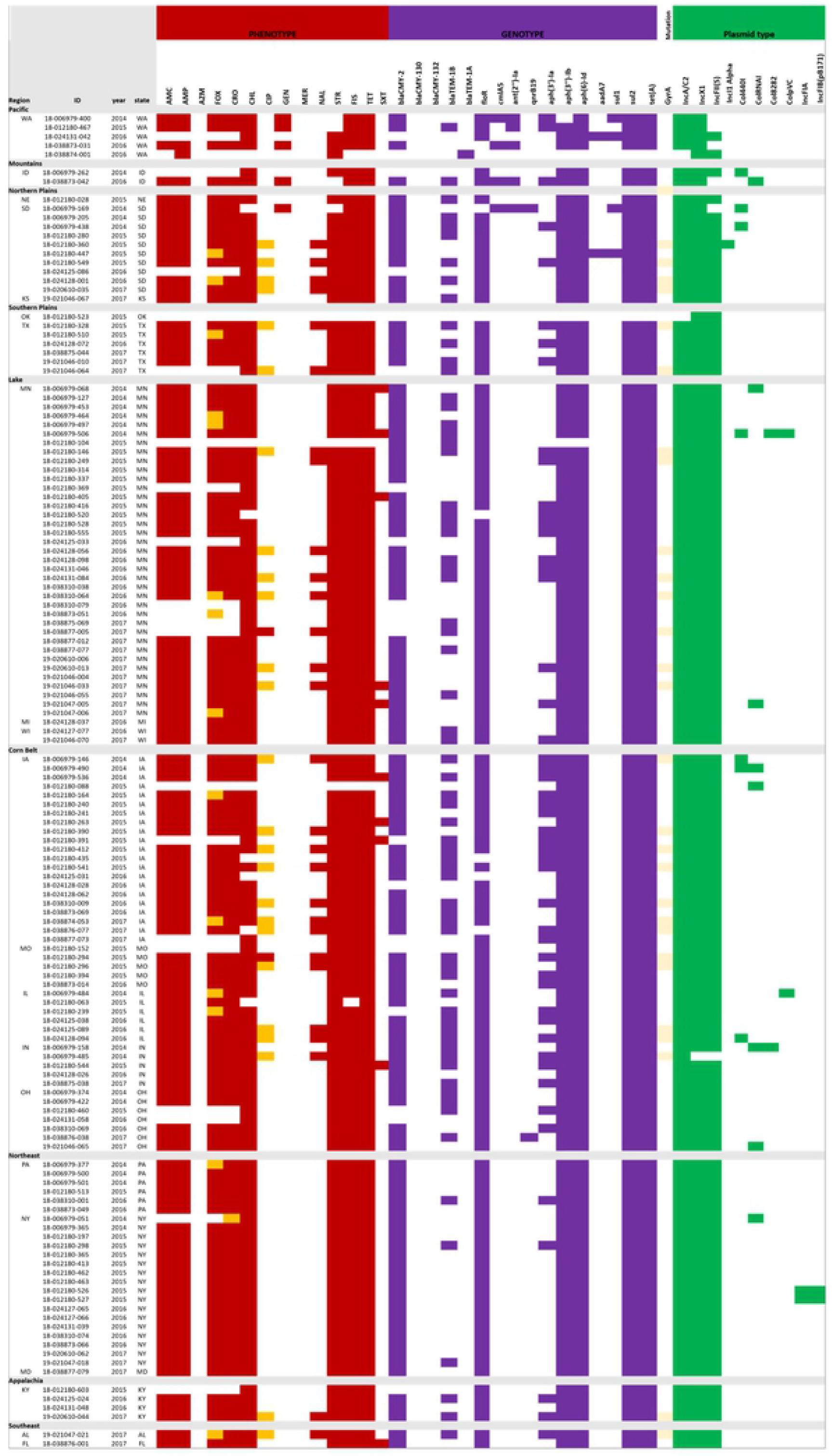
Antimicrobial susceptibility, distribution of ARGs, mutations, and presence of plasmids from the *S*. Dublin isolates in this study by State and Region. Red: resistant, Orange: Intermediate, Violet: presence of gene, Yellow: presence of mutation, Green: presence of plasmid.

(20.00%) isolates exhibited phenotypic resistance with the same gene mutation present.

### Plasmid typing

Ten plasmid types were identified among all isolates (Fig 1): IncA/C2, IncX1, IncFII(S), IncI1 Alfa, Col440I CoIRNAI, Col8282, ColpVC, IncFIA, IncFIB(pB171).

In our study, all isolates contained two to five different plasmid types with the most common ones being: IncX1 (n=139, 99.3%), IncA/C2 (n=138, 98.6%) and IncFII (n= 135, 96.4%).

Multi-drug resistance IncA/C2 plasmid was not found in 2 isolate*s*. One was the pansusceptible isolate, and the second was the isolate resistant only to ampicillin with the presence of the bla-TEM gene. Table 4 shows the antimicrobial susceptibility, distribution of ARGs, mutations, and presence of plasmids from the *S*. Dublin isolates in this study.

Geographic differences were observed for AMR genotypic and phenotypic characteristics (Fig 1). All isolates from New York were susceptible to AZM, CIP, GEN, MER, NAL and TRISUL; and no AMR genes for those antimicrobial drugs were found. All New York isolates were resistant to AMC, AMP, FOX, COX, CHL, STR, FIS and TET with their corresponding genes present.

Annual trends in antimicrobial resistance (2014, 2015, 2016, and 2017) showed no consistent variation overall, but in two states resistance to quinolones showed a slight increase in *gyr*A mutation.

Isolates from MN showed an increase of resistance to quinolones by mutation in *gyr*A, from 0% (n=0 isolates) in 2014, 5.4% (n=2 isolates) in 2015, 8.1% (n=3 isolates) in 2016, and 8.1% (n=3 isolates) in 2017.

In SD, there was also an increase of quinolone resistance. In 2014, the resistance was conferred by qnrB19 for nalidixic acid. In 2015 resistance to nalidixic acid was conferred by mutation in *gyr*A. We also can observe a change of resistance for ciprofloxacin from 0% (n=0 isolates) in 2014, 20% (n=2 isolates) in 2015, 10% (n=1 isolate) in 2016, and 10% (n=1 isolate) in 2017.

### Relationship of antimicrobial susceptibility with antimicrobial genes

Table 3 shows the correlation between phenotype and genotype results. Columns 2 and 3 shows the false positives or false negatives with the phenotype method. A subset of isolates was resistant to trimethoprim sulfamethoxazole, but did not present with any known genetic mechanism of resistance.

**Table 3.**
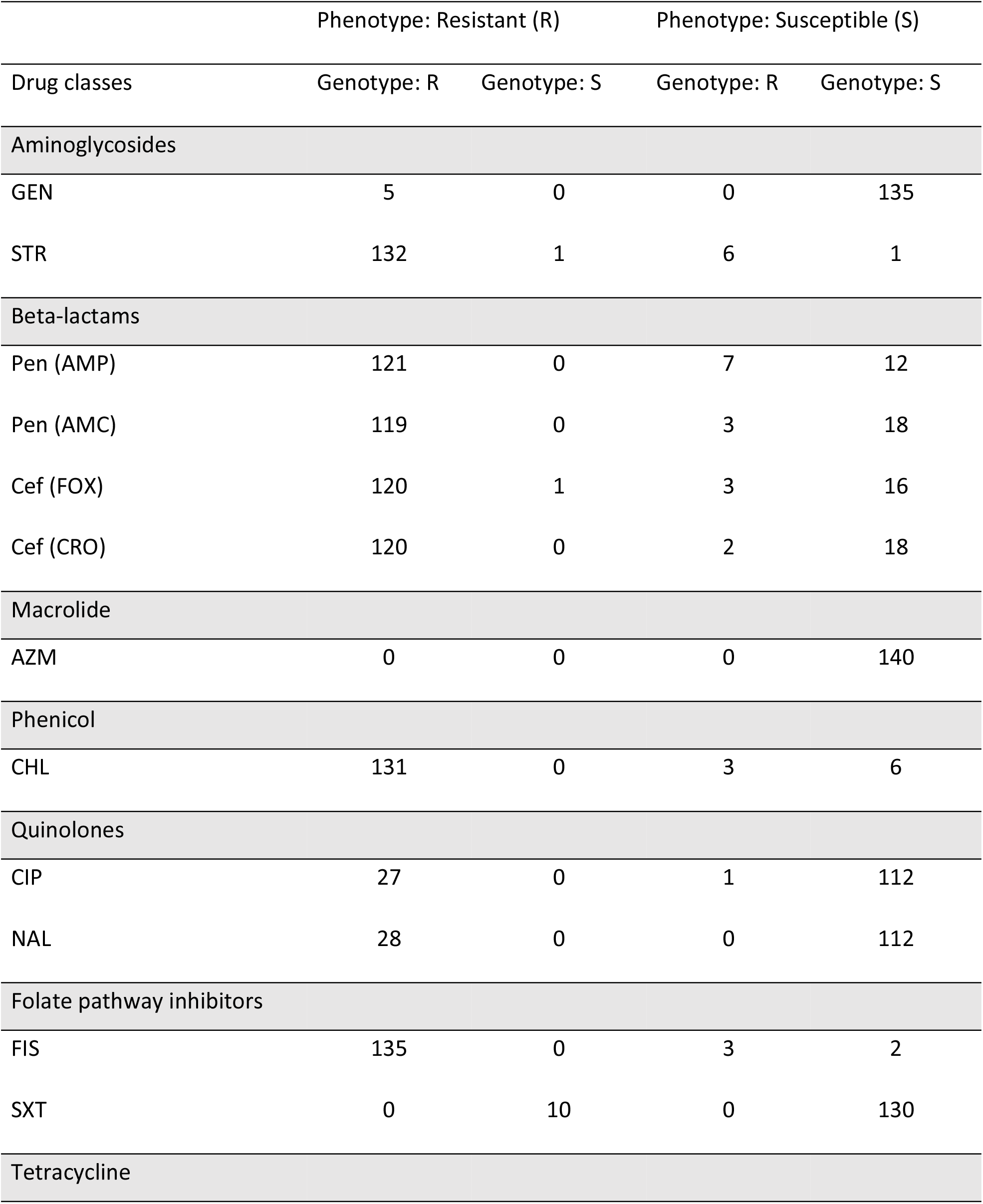
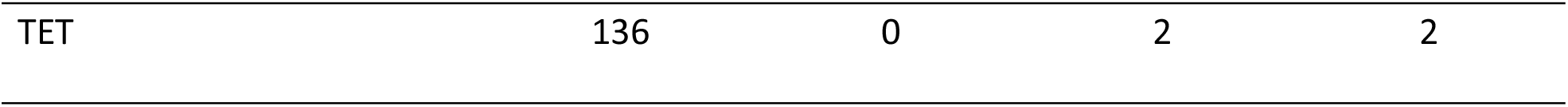
Relationship of antimicrobial susceptibility with antimicrobial genes.

### Phylogenetic relationships

*Salmonella* Dublin isolates are highly clonal. The majority of isolates in this study fall into a single clade with a small number of isolates representing a distinct but closely related branch (Fig 2). All isolates except one were classified into one sequence type (ST), ST10, based on multi-locus sequence type (MLST)analysis from genome sequence*s*.

**Fig 2.**
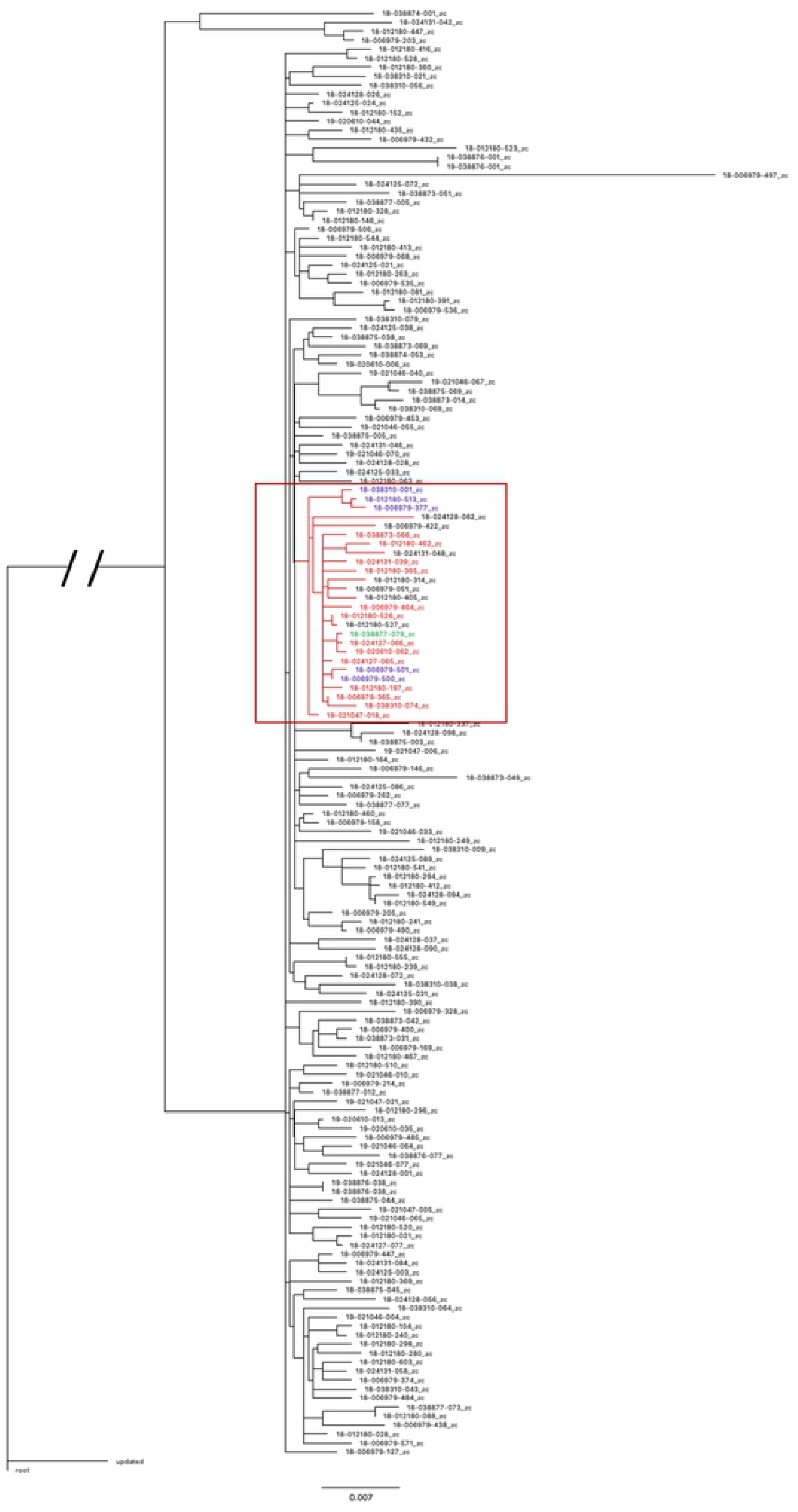
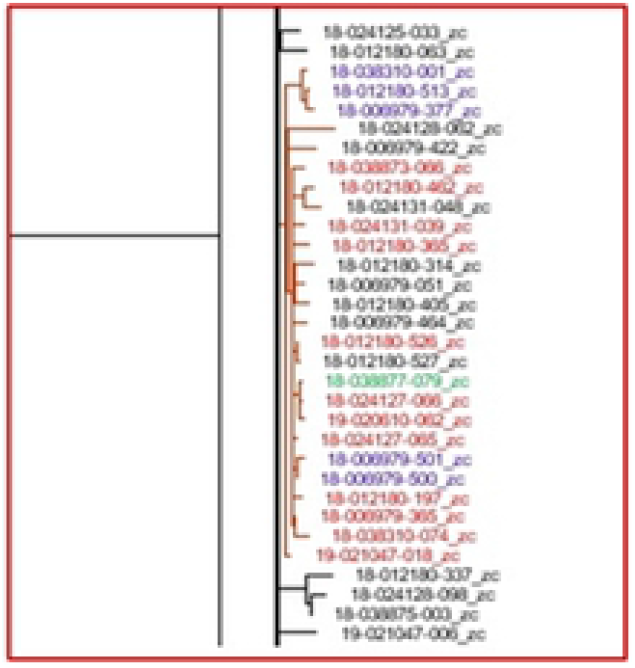
Phylogenetic tree of *S*. Dublin isolates. **a**. All isolates **b**. NY isolates (in red) showing a common relationship on a single branch. Other isolates from northest region are colored (PA isolates in blue and MD isolates in green).

## Discussion

*Salmonella* Dublin has developed into one of the most antimicrobial-resistant serotypes in the United State*s*. A recent study from the FDA [11], showed that among *Salmonella* Dublin isolates recovered from sick cattle and retail meats in Arizona, all except 1 were resistant to >4 antimicrobial classes tested. The results of our study are consistent with the phenotypic resistance profile when the dataset is extended to a nationally distributed sampling.

In our study, the predominant gene conferring beta-lactam resistance was blaCMY-2 (85.7%). Hsu et al. [11], and other studies from different countries also showed that beta-lactam resistance is driven largely by the presence of blaCMY-2 [12]. Another study which compared non-thyphoidal *Salmonella* isolates from humans and retail meat from FDA [13], showed that among beta-lactam-resistant strains, blaCTX-M-14b, blaFOX-6, blaLAP-1, and blaOXA were found only in human isolates and blaSHV-2a and CTX-M-1 were found in retail meat isolates. In the animal isolates from the current study, the predominant gene was blaCMY-2, but blaCMY-130 and CMY-132 were also present. This suggests that the simple presence or absence of resistance in a foodborne pathogen may not be directly correlated to zoonotic risk; and therefore, identification of the genes involved in resistance may be critical to better understand zoonotic infections.

Regarding quinolone resistance, Mangat et al. [12], showed that in ciprofloxacin and nalidixic acid non-susceptible isolates, an altered chromosomal *gyr*A gene was detected. Quinolone resistance-determining region found in our *S*. Dublin isolates (GyrA D87N and GyrA S83F) are similar to those found in other studies [11, 12]. McDermott et al. [13] in the comparison study with *Salmonella* isolated from retail meat and humans, found only quinolone resistance mechanisms (plasmid mediated genes or mutations) in human isolate*s*.

In aminoglycoside resistant isolates, aph(3’’)-Ib and aph(6)-Id genes were also predominantly found in other studies [14, 15]. Additionally, these 2 genes are usually found together in a HI type plasmid [16].

Two AMR genes were found in all isolates using only the Resfinder database. The mdfA gene is a multidrug efflux pump that confers resistance to lipophilic compounds (tetracycline, rifampicin, puromycin) and also confers resistance to chloramphenicol, erythromycin, fluoroquinolones (ciprofloxacin and norfloxacin) and to a much lesser extent, to certain aminoglycosides (neomycin and kanamycin) [17]. The mdfA displays a remarkably broad spectrum of drug recognition. MdfA-associated resistance to aminoglycosides is reproducible, but resistance level is very low and further studies are needed to demonstrate that these drugs are truly exported by mdfA [18]. The level of resistance may be related to the level of expression of MdfA, and the amount of MdfA in the cells is likely not very high. This gene does not appear to correlate to significant resistance in our isolates because this gene was present in all isolates, including the two more susceptible isolates.

The aac(6’)-Iaa gene confers high-level resistance to tobramycin and kanamycin, as well as a significantly increased resistance to amikacin [19]. In the susceptibility determination by microbroth dilution (Sensititre, Thermo Scientific) used in this study, only gentamicin and streptomycin were tested; thus we cannot determine if the gene was expressed.

We found some discordance in the association of genotypic and phenotypic data, as in other studies [20]. The presence of a resistance gene does not necessarily confer phenotypic resistance, and the absence of resistance genes does not unequivocally determine the phenotypic susceptibility. The phenomenon of AMR, then, is not just related to the mere presence or absence of resistance gene*s*. Other mechanisms such as enzyme activation, target modification/protection, regulation of AMR gene expression, or even change in the cell wall charge also play important roles in AMR. We observed that some isolates were resistant to trimethoprim sulfamethoxazole, but they did not present any known resistance gene. This could be due to a promoter, frameshift, or point mutation, for example. Another aspect of a potentiated or combination drug is that resistance may be more multifactorial and less simple to determine genetically.

IncA/C2, IncX1 and IncFII are the most prevalent plasmids present in *S*. Dublin, as in other U.*S*. studies [11, 21]. IncA/C2 is a plasmid associated with MDR, and IncX1-IncFII is associated with virulence. Hsu et al. [11] showed that the IncA/C2 MDR plasmid is commonly present in *S*. Dublin and often carries many ARGs, including blaCMY-2 (65.2%). In our study 85.7% of isolates (n=120) were positive for blaCMY-2 gene, and also carried the IncA/C2 plasmid.

In Canada, Mangat et al. [12] showed that MDR isolates often had an IncA/C2 plasmid. Our results are in concordance with this report: only 2 isolates lacked IncA/C2 gene, one of these did not present any AMR gene, and the other only presented the bla-TEM gene. The carriage of IncA/C2 plasmid was seen as a typical feature of *S*. Dublin isolates from the bovine hosts in China [20].

Virulence plasmid IncX1 was the most prevalent plasmid among our isolates (99.3%); and in a previous study in the U. S. [21], the IncX1 plasmid was detected in all *S*. Dublin genome*s*. This is noteworthy because this plasmid is not detected frequently in other serotypes, including *S*. Typhimurium and *S*. Newport. In the same study [21], Carroll et al. compared salmonellae from dairy cattle and humans in New York State (NY) and Washington State (WA); and geographic differences were observed for AMR genotypic and phenotypic characteristics in isolates from those two states. The IncFII(S) plasmid was more commonly detected in isolates from NY State. This differs from our results where IncFII(S) was the most prevalent plasmid among WA isolates and all NY isolates had IncA/C2, IncFII(S) and IncX1 plasmids in the same proportion. Another interesting result found in the Carroll et al. study was the presence of an allele on a truncated *str*A gene in isolates from the WA State clade which appeared to not confer STR resistance, while still being identified computationally as an STR resistance determinant. Our results showed similar AMR phenotypic and genotypic characteristics in NY isolates; but we could not observe similar characteristics for WA, potentially because our sample size was not very large for this state (only 5 isolates).

Mangat et al. [4] reported a rise in MDR among *S*. Dublin isolates in Canada isolates, and a close relationship with U.S. isolates, similar circulating plasmids and mobile element*s*. Interestingly, the network of MDR isolates was comprised of both human and bovine isolates, whereas the network of susceptible isolates was primarily from human source*s*. A study in China [20] showed that MDR was higher in animal isolates than in human one*s*. The presence of higher rates of MDR in animals as compared to human isolates suggests that antibiotic use in treatment of the severe disease that *S*. Dublin can cause in calves may be a driver for antimicrobial resistance in this serotype; thus management, prevention and intensive supportive treatment may present a more sustainable method for approaching this disease. Use of antimicrobials should be approached cautiously and be informed by antimicrobial susceptibility test information.

All *S*. Dublin isolates are usually identified by MLST as ST10 in most of the studies from different countries [12, 20]. Our results showed ST10 for all isolates except one, in concordance with a study in the United Kingdom [22]. A single sequence type is correlated with a highly conserved serotype such as *S*. Dublin. The vSNP results provide a much higher resolution platform for looking at single SNP changes in the bacteria, creating the potential to look at transmission dynamics and even potentially movement of ARGs on a much finer level. In addition, we could observe that several SNPs were present across multiple isolates that did not otherwise correspond with the phylogenetic relationships; and when we investigated those further, they were mutations associated with antimicrobial resistance. A highly clonal relationship among *S*. Dublin isolates has been reported [11]. The lack of genetic diversity in *S*. Dublin can be largely explained by its unique status as host adapted in cattle. Isolates from New York show an even more conserved population, with 70.6% of isolates of New York origin in the same branch. Some isolates in this branch differ by only 2 SNPs. This may be because of selection pressure or it may represent a stable population of cattle with less movement and transfer of bacteria with other geographic regions. We can observe that NY isolates have a similar ARG profile as well.

While our dataset represents a more geographically diverse sample set of clinical isolates than previous veterinary studies in the United States, expansion of the analysis to include comparison to human- and food-associated isolates from a comparable time period may help to better understand transmission dynamics and the relationship between pathogenic circulating isolates in cattle and those capable of causing significant human disease. In addition, this sample set represents submissions for diagnosis of disease; and therefore, may not be representative of the population of *S*. Dublin strains circulating in normal healthy cattle. A better understanding of the relationship between these populations would help to target intervention strategies at the farm level to *Salmonella* strains that are more likely to cause animal or human disease, allowing for effective targeted intervention.

## Acknowlegements

This project was supported in part by an appointment to the Research Participation Program at the Animal and Plant Health Inspection Service, United States Department of Agriculture, administered by the Oak Ridge Institute for Science and Education through an interagency agreement between the U. S. Department of Energy and USDA APHIS.

## Disclaimer

The findings and conclusions in this publication are those of the authors and should not be construed to represent any official USDA or U.S. Government determination or policy.

## References

1. Harvey RR, Friedman CR, Crim SM, Judd M, Barrett KA, Tolar B, et al. Epidemiology of Salmonella enterica Serotype Dublin Infections among Humans, United States, 1968–2013. Emerg Infect Dis. 2017;23(9):1493–1501.

2. Pl, Fogelman D, Shim SJ, Brunner MA, Lein DH. Salmonella enterica Serotype Dublin Infection: an Emerging Infectious Disease for the Northeastern United States. J Clin Microbiol. 1999; 37(8):2418–2427.

3. Holschbach CL, Peek SF. Salmonella in Dairy Cattle. Vet Clin North Am Food Anim Pract. 2018; 34(1):133–154.

4. Morningstar-Shaw, BR. National Veterinary Services Laboratories (NVSL) Salmonella serotyping report. Proceedings 122^nd^ Annual Meeting US Animal Health Association, Kansas. 2018 Oct 18-24. pp. 280-283. Available from: https://usaha.org/upload/Proceedings/2018_Proceedings_v2_Comb_FINAL.pdf

5. Caudell MA, Dorado-Garcia A, Eckford S, Creese C, Byarugaba DK, Afakye K, et al. Towards a bottom-up understanding of antimicrobial use and resistance on the farm: A knowledge, attitudes, and practices survey across livestock systems in five African countries. Plos One. 2020;15(1):e0220274. doi10.1371/journal.pone.0220274.

6. Feldgarden M, Brover V, Haft DH, Prasad AB, Slotta DJ, Tolstoy I, et al. Validating the AMRFinder Tool and Resistance Gene Database by Using Antimicrobial Resistance Genotype-Phenotype Correlations in a Collection of Isolates. Antimicrob Agents Chemother. 2019;63(11):e00483–19. doi:10.1128/AAC.00483-19. [published correction appears in Antimicrob Agents Chemother. 2020;64(4):e00361-20].

7. Seemann T. Abricate, Github. Available from https://github.com/tseeman/abricate/.

8. Zankari E, Hasman H, Cosentino S, Vestergaard M, Rasmussen S, Lund O, et al. Identification of acquired antimicrobial resistance genes. J Antimicrob Chemother. 2012;67(11):2640–2644.

9. Carattoli A, Zankari E, García-Fernández A, Voldby Larsen M, Lund O, Villa L,et al. In silico detection and typing of plasmids using PlasmidFinder and plasmid multilocus sequence typing. Antimicrob Agents Chemother. 2014;58(7):3895–3903.

10. Ea Zankari, Rosa Allesøe, Katrine G Joensen, Lina M Cavaco, Ole Lund, Frank M Aarestru. PointFinder: a novel web tool for WGS-based detection of antimicrobial resistance associated with chromosomal point mutations in bacterial pathogens. J Antimicrob Chemother. 2017; 72(10):2764–2768.

11. Hsu CH, Li C, Hoffmann M, McDermott P, Abbott J, Ayers S, et al. Comparative Genomic Analysis of Virulence, Antimicrobial Resistance, and Plasmid Profiles of Salmonella Dublin Isolated from Sick Cattle, Retail Beef, and Humans in the United States. Microb Drug Resist. 2019;25(8):1238–1249.

12. Mangat CS, Bekal S, Avery BP, Côté G, Daignault D, Doualla-Bell F, et al. Genomic investigation of the emergence of invasive multidrug resistant Salmonella Dublin in humans and animals in Canada. Antimicrob Agents Chemother. 2019;63(6):e00108–19. doi: 10.1128/AAC.00108-19.

13. McDermontt PF, Tyson GH, Kabera C, Chen Y, Li C, Folster JP, et al. Whole-Genome Sequencing of fetecting Antimicrobial Resistance in Nontyphoidal Salmonella. Antimicrob Agents Chemother. 2016; 60(9):5515–5520.

14. Yasit M, Farman M, Shah MW, Jiman-Fatani AA, Othman NA, Almasaudi SB, et al. Genomic and antimicrobial resistance genes diversity in multidrug-resistant CTX-M-positive isolates of Escherichia coli at a health care facility in Jeddah. J Infect Public Health. 2020;(13):94–100.

15. Cohen E, Davidovich M, Rokney A, Valinsky L, Rahav G, Gal-Mor O. Emerge of new variant of antibiotic resistance islands among multidrug-resistant Salmonella enterica in poultry. Environ Microbiol. 2019;22(1):413–432.

16. MacMillan EA, Gupta SK, Williams LE, Jove T, Hiott LM, Woodley TA, et al. Antimicrobial Resistance genes, cassettes, and plasmids present in Salmonella enterica associated with United States food animals. Front Microbiol. 2019;10:832. doi:10.3389/fmicb.2019.00832.

17. Adzitey F, Assoah-Peprah P, Ayum TG. Whole-genome sequencing of Eschrichia coli isolated from contaminated meat samples collected from the Northern region of Ghana reveals the presence of multiple antimicrobial resistance genes. J Glob Antimicrob Resist. 2019;18:179–182.

18. Edgar R, Bibi E. MdfA, an Escherichia Resistance Protein with an Extraordinarily Broad Spectrum of Drug Recognition. J Bacteriol. 1997;179(7):2274–2280.

19. Salispante S, Hall B. Determining the limits of the evolutionary potential of an antibiotic resistance gene. Mol Biol Evol, 2003;20(4):653–659.

20. Paudyal N, Pan H, Elbediwi M, Zhou X, Peng X,Li X, et al. Characterization of Salmonella Dublin isolated from bovine and human hosts. BMC Microbiol. 2019;19(1):226. doi:10.1186/s12866-019-1598-0.

21. Carroll LM, Wiedmann M, den Bakker H, Siler J. Whole-Genome Sequencing of Drug-Resistant Salmonella enterica Isolates from Dairy Cattle and Humans in New York and Washington States Reveals Source and Geographic Associations. Appl Environ Microbiol. 2017;83(12):e00140–17. doi:10.1128/AEM.00140-17.

22. Mohammed M, Le Hello S, Leekitcharoenphon P, Hendriksen R. The invasome of Salmonella Dublin as revealed by whole genome sequencing. The invasome of Salmonella Dublin as revealed by whole genome sequencing. BMC Infect Dis. 2017;17(1):544. doi:10.1186/s12879-017-2628-x.

